# Contrast-free Super-resolution Doppler (CS Doppler) based on Deep Generative Neural Networks

**DOI:** 10.1101/2022.09.29.510188

**Authors:** Qi You, Matthew R. Lowerison, YiRang Shin, Xi Chen, Nathiya Vaithiyalingam Chandra Sekaran, Zhijie Dong, Daniel A. Llano, Mark A. Anastasio, Pengfei Song

## Abstract

Super-resolution ultrasound microvessel imaging based on ultrasound localization microscopy (ULM) is an emerging imaging modality that is capable of resolving micron-scaled vessels deep into tissue. In practice, ULM is limited by the need for contrast injection, long data acquisition, and computationally expensive post-processing times. In this study, we present a contrast-free super-resolution Doppler (CS Doppler) technique that uses deep generative networks to achieve super-resolution with short data acquisition. The training dataset is comprised of spatiotemporal ultrafast ultrasound signals acquired from *in vivo* mouse brains, while the testing dataset includes *in vivo* mouse brain, chicken embryo chorioallantoic membrane (CAM), and healthy human subjects. The *in vivo* mouse imaging studies demonstrate that CS Doppler could achieve an approximate 2-fold improvement in spatial resolution when compared with conventional power Doppler. In addition, the microvascular images generated by CS Doppler showed good agreement with the corresponding ULM images as indicated by a structural similarity index of 0.7837 and a peak signal-to-noise ratio of 25.52. Moreover, CS Doppler was able to preserve the temporal profile of the blood flow (e.g., pulsatility) that is similar to conventional power Doppler. Finally, the generalizability of CS Doppler was demonstrated on testing data of different tissues using different imaging settings. The fast inference time of the proposed deep generative network also allows CS Doppler to be implemented for real-time imaging. These features of CS Doppler offer a practical, fast, and robust microvascular imaging solution for many preclinical and clinical applications of Doppler ultrasound.

## I. Introduction

Super-resolution ultrasound microvessel imaging is a rapidly growing field. Early studies conducted by Viessmann *et al*. [1], Desailly *et al*. [2-3], and O’Reilly *et al*. [4] showed that one can break the resolution limit of acoustic waves by localizing microbubbles in the blood stream. The seminal papers by Errico *et al*. [5] and Christensen-Jeffries *et al*. [6] catalyzed the growth of the field, leading to many subsequent reports with successful *in vivo* applications [7-16] and technical advancements of super-resolution ultrasound imaging [17-32].

At present, most super-resolution ultrasound imaging techniques rely on microbubble (MB) localization, e.g., ultrasound localization microscopy (ULM) [6]. However, these techniques are challenged by the competing goals of minimizing data acquisition time and reconstructing microvessel images that are complete [33-35]. For instance, to reconstruct the 2D cross-section of a vessel with a diameter of 100 µm, one needs at least 50 independent microbubble events occurring at unique spatial locations (assuming a uniform MB diameter of 2 µm). However, in practice, this is a challenging task because MB events are dictated by local MB concentration and blood flow and therefore entirely stochastic. Consequently, a long data acquisition time is typically required to accumulate adequate MB signals to fully reconstruct the tissue vasculature. As such, the temporal resolution of ULM is inherently limited by the nature of MB flow in the blood stream and the MB localization process. Another pragmatic challenge of ULM is that a relatively stable MB concentration is necessary to ensure optimal image quality. However, MB concentration is difficult to control in practice, especially with bolus injections that are commonly used in preclinical and clinical practices. Therefore, the objective of this study was to explore an alternative solution for super-resolution microvascular imaging without the need of contrast MBs or the localization process.

In the past decade, a substantial body of literature has reported the combination of long Doppler ensembles with advanced clutter filters (e.g., the ones based on singular value decomposition (SVD)) to increase Doppler sensitivity to small vessels [36-38]. In addition to improving Doppler sensitivity (e.g., SNR), several methods were recently proposed to improve the spatial resolution of contrast-free power Doppler [39-42]. For example, Bar-Zion *et al*. [39] used hundreds of narrowband filters to slice the Doppler signal in the temporal dimension and performed sparsity-based reconstruction on individual signal slices. This technique demonstrates improved spatial resolution but still requires relatively long data acquisition to be effective; Lok *et al*. [40] used deconvolution with total variation regularization to improve the spatial resolution of contrast-free microvascular imaging. This technique does not require long data acquisition time and is computationally efficient. However, the deconvolution technique requires *a priori* knowledge of the point spread function (PSF) and noise distribution of the ultrasound system, which are difficult to obtain in practice. Jensen et al. [41] and Park et al. [42] reconstructed microvessel images by localizing speckle patterns of red blood cells (RBCs). However, the accuracy and efficiency RBC localization may be compromised by the dense distribution of RBCs in the blood flow.

Super-resolving the tissue microvasculature using backscattering signals from RBCs is challenging because RBC signals are much weaker and denser than MB signals. Consequently, using native RBCs for super-resolution imaging becomes a challenging problem. Inspired by the recent advances of super-resolution image reconstruction based on deep generative networks in various biomedical imaging modalities [43-47], in this work we propose to use deep learning to achieve contrast-free super-resolution microvessel imaging based on RBCs for ultrasound. Our hypothesis is that deep generative networks are capable of 1) learning the distributions of contrast-free ultrasound data and ULM images; 2) translating contrast-free ultrasound data into super-resolved microvascular images by minimizing the distance between these two distributions. However, different from conventional deep learning-based super-resolution image reconstruction techniques where pairs of stationary low-resolution and high-resolution images are used for training, we propose to use spatiotemporal ultrasound data for training. The motivation for using spatiotemporal data comes from observations that single image super-resolution reconstruction techniques are susceptible to hallucinations [48-49], which is a major concern for biomedical imaging. Our hypothesis is based on the super-resolution optical fluctuation imaging technique (SOFI) [50] where it was demonstrated that imaging resolution can be improved by leveraging the high order statistics of the fluctuating signal in the temporal dimension. A similar technique [21] has also been applied in the ultrasound field which achieved an enhanced spatiotemporal resolution for microvascular imaging. In this paper, we designed a U-Net-structured deep generative network that generates super-resolution microvessel images from spatiotemporal contrast-free ultrasound signal. A variety of loss functions including the conventional L1 loss, the feature space loss, and the adversarial loss were integrated and investigated in the proposed generative network.

The remainder of the paper is organized as follows. In Section II, we present the mathematic model of the proposed super-resolution microvascular reconstruction technique, followed by the deep neural network training strategy. In Section III, the results from the *in vivo* mouse brain study, *in vivo* chicken embryo chorioallantoic membrane (CAM) study and *in vivo* human study are presented. In Section IV and V, we finalize the paper with discussion and conclusions.

## II. Method

### A. Principles of CS Doppler

The three-dimensional ultrasound signal subject to the SVD clutter filter [23] can be expressed as:

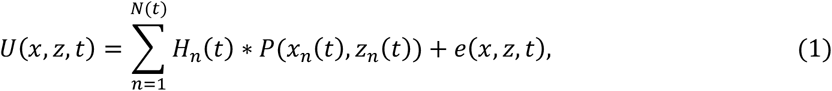

where *U*(*x, z, t*) represents the spatiotemporal ultrasound signal (in 2D imaging) with (*x, z, t*) denoting lateral, axial and temporal dimension, respectively; *n* is an index for the scatterers in the blood flow (e.g., index of MBs in contrast-enhanced ultrasound or index of RBCs in contrast-free imaging); *N(t)* is the total number of scatterers at time *t*; *H*_*n*_(*t*) represents of the impulse response of the *n*^*th*^ scatterer at time *t*; *P*_*n*_(*x*_*n*_(*t*), *z*_*n*_(*t*)) represents the ultrasound imaging system PSF at location *x*_*n*_(*t*), *z*_*n*_(*t*), where *x*_*n*_(*t*) and *z*_*n*_(*t*) are the lateral and axial locations of the *n*^*th*^ scatterer at time *t*; and *e(x, z, t)* represents the electronic noise at time *t*.

In localization-based super-resolution microvascular imaging, the task is to recover the locations of the scatterers, *(x*_*n*_(*t*), *z*_*n*_(*t*)), followed by accumulation of locations to reconstruct super-resolved microvessel images S(*x, z*):

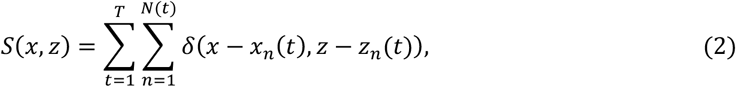

where *T* represents the total imaging time; *δ* represents the Dirac delta function. In ULM, the task of obtaining *S*(*x, z*) is achieved by various MB localization algorithms [5-7,23].

For the proposed contrast-free super-resolution reconstruction, we use a deep neural network *f* to approximate *S*(*x, z*):

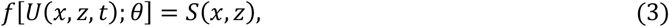

where *θ* represents the parameters of the neural network, which are estimated by minimizing a loss function *L*. The minimization of the loss function is calculated using the *M* training data, where *M* denotes the number of training data pairs that include the spatiotemporal ultrasound signal *U*(*x, z, t*) and its corresponding ULM image:

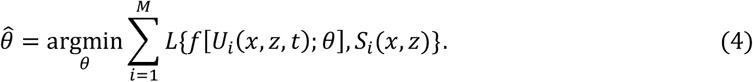

After training, the network *f* learns the translation from the original ultrasound signal *U*(*x, z, t*) to the super-resolved vessel structure *S*(*x, z*). *L*{·} represents the objective function for calculating the training loss.

In this study, we have adopted several complementary loss functions including the L1 loss, the feature space loss [51] and the adversarial loss [52]. These loss terms have been widely used in a variety of super-resolution image reconstruction applications including ULM, magnetic resonance imaging (MRI), optical and photoacoustic microscopy, and computer vision [22, 43-47]. The L1 loss promotes the underlying spatial sparsity within the microvascular structure [22] but struggles to recover high-frequency details [45], which can be complemented by the adversarial loss that minimizes the distance of the low-dimensional manifold between the two data distributions [45-46]. We also incorporated the feature space loss that minimizes the Euclidean distance between the feature maps extracted from the VGG19 network [51], which improves the perceptual quality of the recovered super-resolution images [45-46,53]. In this study, three different objective functions were used:

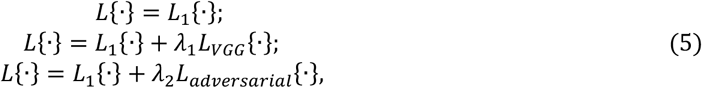

where *L*_*1*_{·}, *L*_*VGG*_{·} and *L*_*adversarial*_{·} represent the L1 loss, the feature space loss and the adversarial loss respectively; λ_*1*_ and λ_*2*_ are the regularization parameters, whose value were empirically determined as λ_*1*_ = 0.001 and λ_*2*_ = 0.01 to allow CS Doppler to achieve the visually optimal reconstructed images. The performance of different loss terms was systematically tested for the proposed CS Doppler technique.

### B. Deep Neural Network Design

Figure 1 shows the schematic of the designed deep neural network for CS Doppler. The input to the neural network is the spatiotemporal ultrasound signal *U*(*x, z, t*) after clutter filtering. The generator network represents the deep neural network function *f* used for reconstructing the CS Doppler image from the input ultrasound signal. The corresponding ULM image represents the ground truth and was used for calculating the training loss using the objective function *L*. The training loss was then backpropagated to the generator to adjust the network parameters using the Adam gradient descent algorithm [54].

**Fig. 1.**
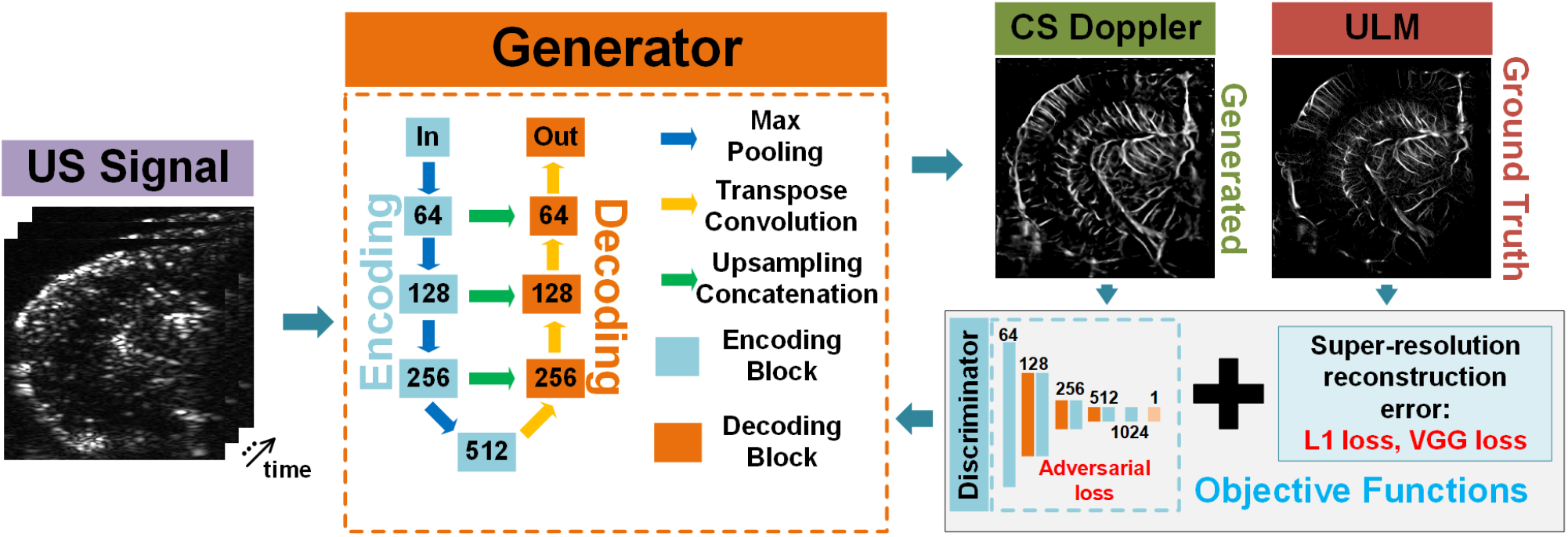
Architecture of the proposed deep generative network. The generator takes the spatiotemporal ultrasound signal as input and outputs an estimate of the CS Doppler image. The generated CS Doppler image is compared with the ULM ground truth image to calculate the training loss through the objective function, which was then backpropagated to the generator. The numbers marked in the generator and discriminator represent the number of channels for the convolutional layers within the respective network.

The generator was designed using a U-Net encoding-decoding architecture. In the encoding part, each encoding block consists of two 3 × 3 convolutional layers with batch normalization (BN) and rectified linear unit (ReLU) activations, followed by a 2 × 2 max pooling layer to down-sample the image. In the decoding part, each decoding block includes a 2 × 2 transpose convolutional layer with a stride of 2 and two 3 ×3 convolutional layers with BN and ReLU. An additional 2 × 2 bilinear up-sampling layer was added after each decoding block to increase the image size. The concatenation between encoding and decoding blocks also includes a bilinear up-sampling layer to match the size of images between encoding and decoding blocks.

To calculate the adversarial loss, a discriminator network was used to output a probability for distinguishing the generated CS Doppler image and the ULM ground truth. The discriminator network [45] includes one input block, one output block, and three down-sampling blocks. The input and output blocks are composed of a 3 ×3 convolutional layer. The down-sampling blocks include two 3 × 3 convolutional layers, where the latter one uses a stride of 2 to down-sample the image. All the convolutional layers in the discriminator used leaky ReLU activation with a negative slope of 0.2. The discriminator and the generator were trained simultaneously by minimizing the approximation of the Earth-Mover (EM) distance [55].

### C. Training and Testing Methods

*In vivo* data from mouse brain was used in this study for training and testing. All mouse brain data were acquired using a Verasonics Vantage 256 System (Verasonics, Kirkland, WA, USA) and a L35-16vX transducer (Verasonics, Kirkland, WA, USA). In addition to the mouse brain, chicken embryo CAM data were also acquired with the Verasonics system and the L35-16vX transducer. Imaging was performed using plane-wave compounding with steering angles from −4° to 4° with a step size of 1°. The post-compounding frame rate was 1,000 Hz. Electrocardiogram (ECG) signals of the mouse were acquired using an iWorx (Dover, NH, USA) IA-100B single channel biopotential amplifier with C-MXLR-PN3 platinum needle electrodes inserted into the limbs of the mouse. The details of the animal procedure were provided in our previous studies [12,34].

For the *in vivo* human data, the kidney and liver from a healthy volunteer were used in this study. The data is identical to the ones used in [38], which was acquired using a L11-4v transducer (Verasonics Inc., Kirkland, WA) operating at 5 MHz with plane-wave compounding from −9° to 9° with a step size of 2° at a post-compounding frame rate of 500 Hz. The human data were only used for testing.

The ultrasound signal was processed using the SVD clutter filter [38] and the flow separation filters [25], then input into the deep generative network to generate the CS Doppler images. The ULM images were reconstructed using a Kalman filter based localization and tracking algorithm [23-24]. All the animal experiments in this study were approved by the Institutional Animal Care and Use Committee (IACUC) at the University of Illinois Urbana-Champaign.

The design of the training, validation and testing dataset was listed in Table I. We acquired the ultrasound data from 51 different image planes in 17 mouse brains (3 imaging planes from each mouse). For robust network training with large datasets, both contrast-enhanced and contrast-free ultrasound data were used for training. This study design was justified by the similar flowing pattern and data characteristics of the RBC and MB signals in the blood stream. Specifically, nine blocks of 400 frames of contrast-enhanced ultrasound data and one block of 400 frames of contrast-free ultrasound data were acquired for each imaging plane as the network input. One ULM image reconstructed using 32000 frames was used as the ground truth. The training dataset was augmented by a factor of 8 using the flow separation filter [25], where 10% of the contrast-enhanced and contrast-free training dataset were randomly selected as the validation dataset ahead of the training process. The validation dataset was used for a learning rate scheduler which initialized with a learning rate of 0.001 and decayed by a factor of 0.1 when the validation loss stopped improving for 5 epochs. The number of epochs used for training was set to 50. For testing, we acquired ultrasound data from 6 imaging planes from 2 additional mice (3 imaging planes from each mouse) whose data were never used for training. One block of 400 frames of contrast-enhanced and contrast-free ultrasound data from each imaging plane was used as the network input and one ULM image reconstructed using 32000 frames was used as the ground truth. Finally, 400 frames of contrast-free ultrasound data from a chicken embryo CAM, a human liver, and a human kidney were acquired to test the generalization ability of the proposed network.

**Table I.** Design of the training, validation and testing dataset.

|  | **Data Acquisition** | **Network Input** | **Ground Truth** |
| --- | --- | --- | --- |
| **Mouse Brain** | **Training:** 51 imaging planes in 17 mice (3 imaging planes each mouse) | 10 blocks of 400 frames each imaging plane | 1 ULM accumulating 32000 frames each imaging plane |
|  | **Validation:** 51 imaging planes in 17 mice (3 imaging planes each mouse) | Randomly selected 10% of the training dataset | 1 ULM accumulating 32000 frames each imaging plane |
|  | **Testing:** 6 imaging planes in 2 mice (3 imaging planes each mouse) | 2 block of 400 frames each imaging plane | 1 ULM accumulating 32000 frames each imaging plane |
| **Chicken Embryo CAM/Human liver/Human Kidney** | **Testing:** 1 imaging plane | 1 block of 400 frames each imaging plane |  |

### D. CS Doppler Performance Evaluation

The imaging performance of the proposed CS Doppler technique was assessed by calculating the ensemble-averaged estimates of the peak signal-to-noise ratio (PSNR) and the structural similarity index measure (SSIM) [54] using all the data in the testing dataset. The PSNR and SSIM are defined by [56]

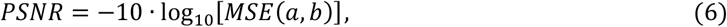

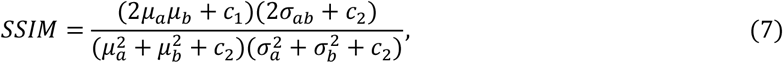

where *a* and *b* represent the normalized generated image and the ground truth, respectively; *MSE(*·) represents the mean squared error; *µ*_*a*_ and *µ*_*b*_ represent the average intensity of *a* and *b* after applying a 3 ×3 Gaussian window; *σ*_*a*_, *σ*_*b*_ and *σ*_*ab*_ represent the variance of *a*, b and the covariance of *a*, b after applying the 3 ×3 Gaussian window. PSNR and SSIM were used to evaluate the pixel-wise and structural differences between the generated image and the ground truth.

In addition to PSNR and SSIM, we also used full width at half maximum (FWHM) and MSE of the normalized intensity profile (MSE-NIP) as the specific measurements for the individual vessels. The FWHM was measured on the cross-sectional profile of local vessels to evaluate the resolution improvement of CS Doppler. MSE-NIP was measured on the longitudinal profile of the local vessels to evaluate the consistency of the vessel intensity distribution between the CS Doppler and ULM.

## III. Result

### A. Significance of Temporal Information

Figure 2 compares the CS Doppler results using different implementations of spatiotemporal ultrasound signal as input. All CS Doppler images and corresponding power Doppler images in Fig. 2 were reconstructed using the contrast-free data acquired from a mouse brain selected from the testing dataset. Three different input strategies were used and compared: 1) adjusting the number of input frames for both network training and testing; 2) truncating the number of input frames only in testing (i.e., the network was trained using the full 400 frames of ultrasound signal, the test data input ultrasound signal was truncated to a subset of frames, and then linearly interpolated in the temporal dimension to 400 frames to match the network architecture weightings); 3) using the single 2D power Doppler image after accumulating the ultrasound signal as the network input for both training and testing.

**Fig. 2.**
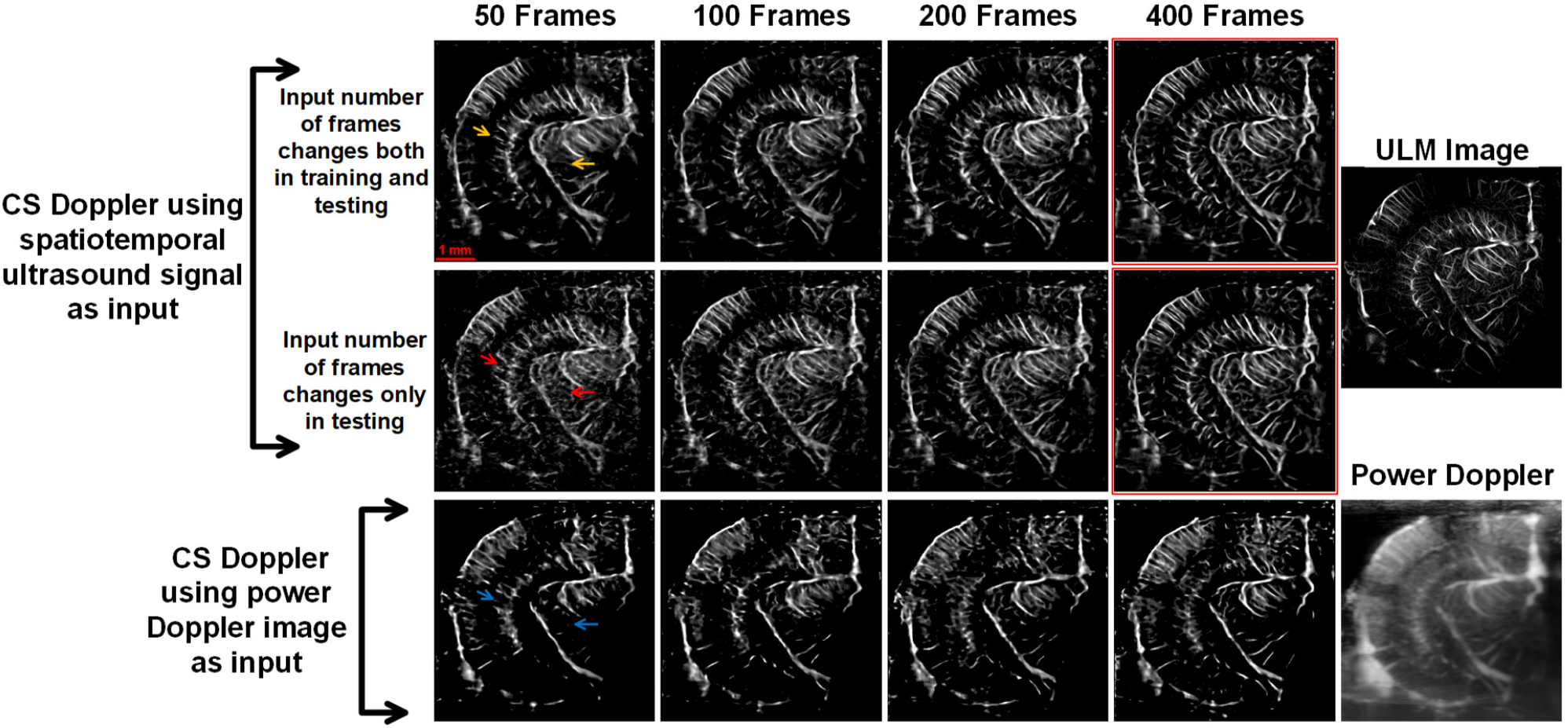
CS Doppler using different ways to input ultrasound signal: input number of frames changing both in the network training and testing (first row); input number of frames changing only in the network testing (second row); inputting power Doppler image after accumulating the spatiotemporal ultrasound signal (third row). The corresponding ULM image and power Doppler image of the same image plane were shown on the right side.

It can be seen in Fig. 2 (b) that the network cannot recover some of the small vessel structures (indicated by the blue arrows) when using single 2D power Doppler images as input (i.e., without temporal information). Moreover, hallucination structures (indicated by the red arrows) were created with low number of input frames and a discrepancy between the number of training and testing input frames. In comparison, when using the same number of frames in both training and testing, CS Doppler can more robustly reconstruct the microvessel structures with less hallucinations (indicated by the orange arrows). As the number of input frames increases, more and more temporal blood flow information was included in training and testing, which leads to improved CS Doppler image quality with more detailed cerebrovasculature (see images marked by red outline). Results in Fig. 2 reveal that the inclusion of temporal information is essential for the network to differentiate and resolve microvessels with different flowing characteristics, which are otherwise indiscernible with single 2D power Doppler images as input.

### B. Evaluation of the performance of CS Doppler

Figure 3 shows the comparison among conventional Doppler (power Doppler and deconvolution-based Doppler), CS Doppler using different loss functions, and the corresponding ULM image. The conventional Doppler images and CS Doppler images were reconstructed using the same 0.4s (400 frames) of contrast-free data from the testing dataset. The corresponding ULM image was reconstructed using 32s of ultrasound data (corresponding to 32000 frames) acquired from the same image plane after injecting contrast MBs. The results clearly indicate that CS Doppler substantially improved the spatial resolution over conventional Doppler including the deconvolution-based resolution enhancement approach. In the cross-sectional vessel profile measurement (Fig. 3e, top panel), CS Doppler improved the FWHM of a penetrating cortical vessel from 50μm to 27μm, which is similar to the 22μm FWHM measured from the ULM image.

**Fig. 3.**
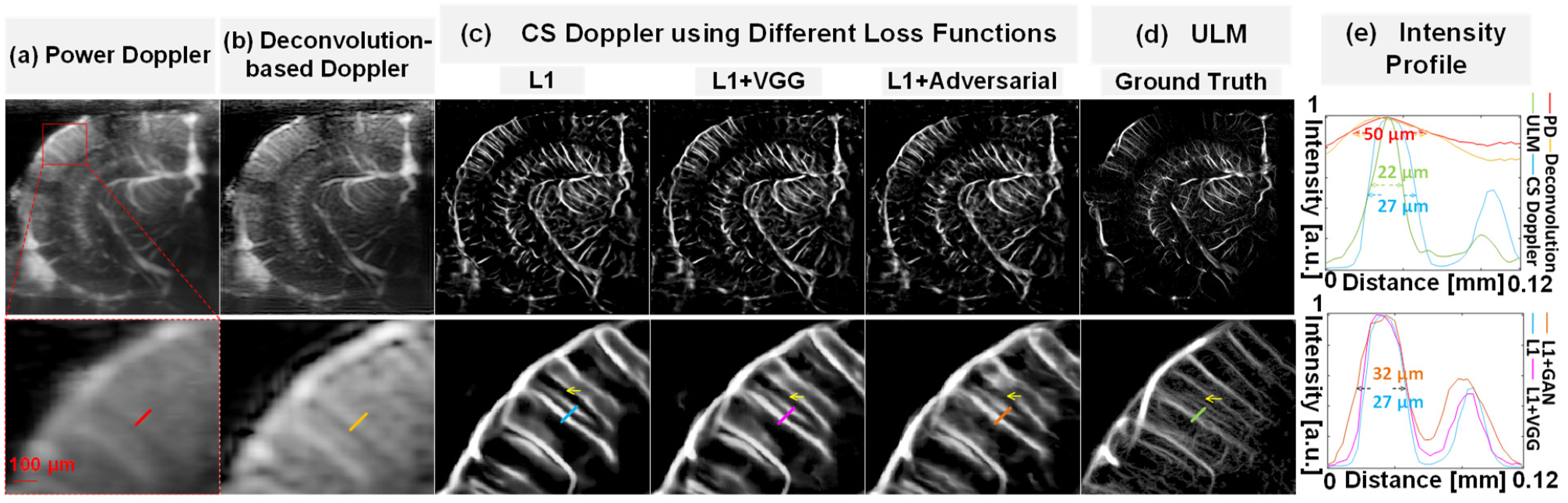
(a) Conventional power Doppler image accumulating 400 frames of contrast-free ultrasound data (b) Deconvolution-based Doppler image reconstructed using 400 frames of contrast-free ultrasound data. (c) CS Doppler images using different loss functions reconstructed using 400 frames of contrast-free ultrasound data. (d) ULM image reconstructed using 32000 frames of contrast-enhanced ultrasound data. (e) The intensity profile of the small penetrating cortical vessels marked by the short line segments with different colors. All the intensity profiles were normalized in the linear scale. All the zoomed-in local vessel images (bottom row) correspond to the same region marked by the red box in the power Doppler image (a).

Table II summarizes the quantitative metrics used to evaluate imaging performance among various techniques. The ensemble-averaged estimates of SSIM and PSNR were evaluated using all the data in the testing dataset. The MSE-NIP and FWHM ratio were evaluated by selecting 10 local vessels in the mouse brain images from the testing dataset. The MSE-NIP was measured on the longitudinal profiles of manually selected vessels between CS Doppler and ULM images. The FWHM ratio was the measured by dividing the FWHM of the cross-sectional profiles of manually selected vessels between power Doppler and CS Doppler. Two examples of such selected local vessel profiles are shown in Figure 4. All the quantitative metrics were measured for the mean and the standard deviation (indicated by the number after ± symbol) across all the samples used for evaluation as shown in Table II. Among all the CS Doppler images, results using L1 with VGG loss and L1 with adversarial loss show more small vessel structures especially for regions that are in between the parallel penetrating vessels (Fig. 3c). However, images generated with L1 loss show the highest SSIM and MSE-NIP measurements (Table II), which indicate the highest consistency with the ULM ground truth. In addition, the vessel profile measured from CS Doppler images generated by the L1 loss (Fig. 3e, bottom panel) was also closest to ULM. Collectively, these results indicate that CS Doppler using L1 loss achieved the best performance in terms of SSIM and MSE-NIP but had the worst performance in terms of PSNR and FWHM ratio. As shown in Fig. 3, one reason for the poor PSNR is that the L1 loss missed small vessel structures (indicated by the yellow arrows in Fig. 3) that were recovered by using L1 loss combined with VGG and L1 combined with adversarial loss. However, the low SSIM and MSE-NIP scores from L1 loss combined with VGG or adversarial loss indicate that the recovered small structures may not be valid (i.e., hallucinations) when comparing to ULM. These results imply that due to the lack of contrast enhancement and the subsequent absence of distinct point scatterers like MBs, the small vessel recovery may be fundamentally limited in CS Doppler.

**Table II.** Evaluation metrics of CS Doppler using different loss functions.

| Metrics<br>Loss | SSIM | PSNR (dB) | NIP-MSE | FWHM Ratio<br>(Power Doppler/CS Doppler) |
| --- | --- | --- | --- | --- |
| **L1** | **0.784±0.095** | 24.645±1.650 | **0.061±0.039** | 2.277±0.593 |
| **L1+VGG** | 0.779±0.090 | 25.105±1.819 | 0.120±0.057 | 2.327±0.813 |
| **L1+Adversarial** | 0.783±0.090 | **25.518±1.840** | 0.067±0.024 | **2.378±0.836** |

**Fig. 4.**
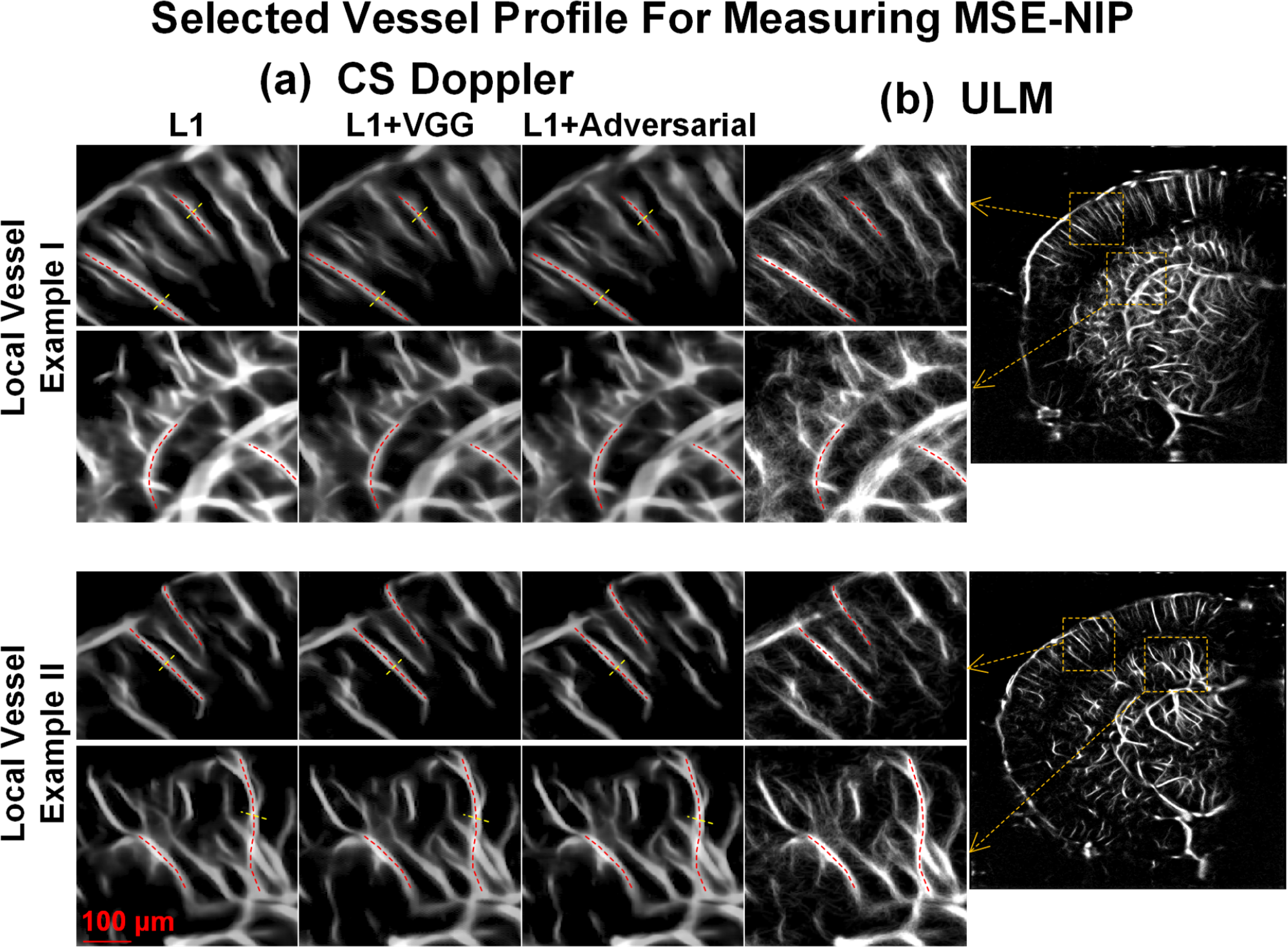
Two examples of the selected vessel profiles for measuring the MSE-NIP (red curves) and FWHM ratio (yellow curves) in Table II. (a) CS Doppler using different loss functions. (b) ULM image of the same imaging plane. All the profiles were manually selected along the longitudinal or cross-sectional direction of the vessels.

### C. Evaluation of the temporal resolution of CS Doppler

Figure 5 shows the comparison of CS Doppler and conventional power Doppler using different number of frames as input and the temporal intensity changes of a local vessel. The conventional Doppler images and CS Doppler images were reconstructed using the same contrast-free data from the testing dataset. For CS Doppler using less than 400 frames, the number of input frames changed both in the network training and testing. Figure 5 (c) also showcases the ability of preserving temporal information (e.g., pulsatility of the blood flow) for CS Doppler. ULM image reconstructed using 32s of data acquisition (32000 frames) was used as the ground truth. To extract the temporal blood flow signal, power Doppler and CS Doppler were reconstructed using 0.2s (200 frames) temporal sliding window with a step size of 0.05s (50 frames). Supplementary video 1 demonstrates the dynamic flow signal revealed by CS Doppler and power Doppler. Based on results from above, only L1 loss was used for CS Doppler. As shown in Fig. 5(c), the blood flow pulsatility measured from CS Doppler showed good agreement with that from conventional Doppler. In addition, both pulsatility measurements showed good agreement with the ECG signal, with the peak of the blood flow occurring after the peak R-wave which indicates peak systole. These results indicate that CS Doppler has similar temporal resolution as conventional power Doppler and is capable of preserving the important temporal information of blood flow, which is significant for applications such as functional brain imaging.

**Fig. 5.**
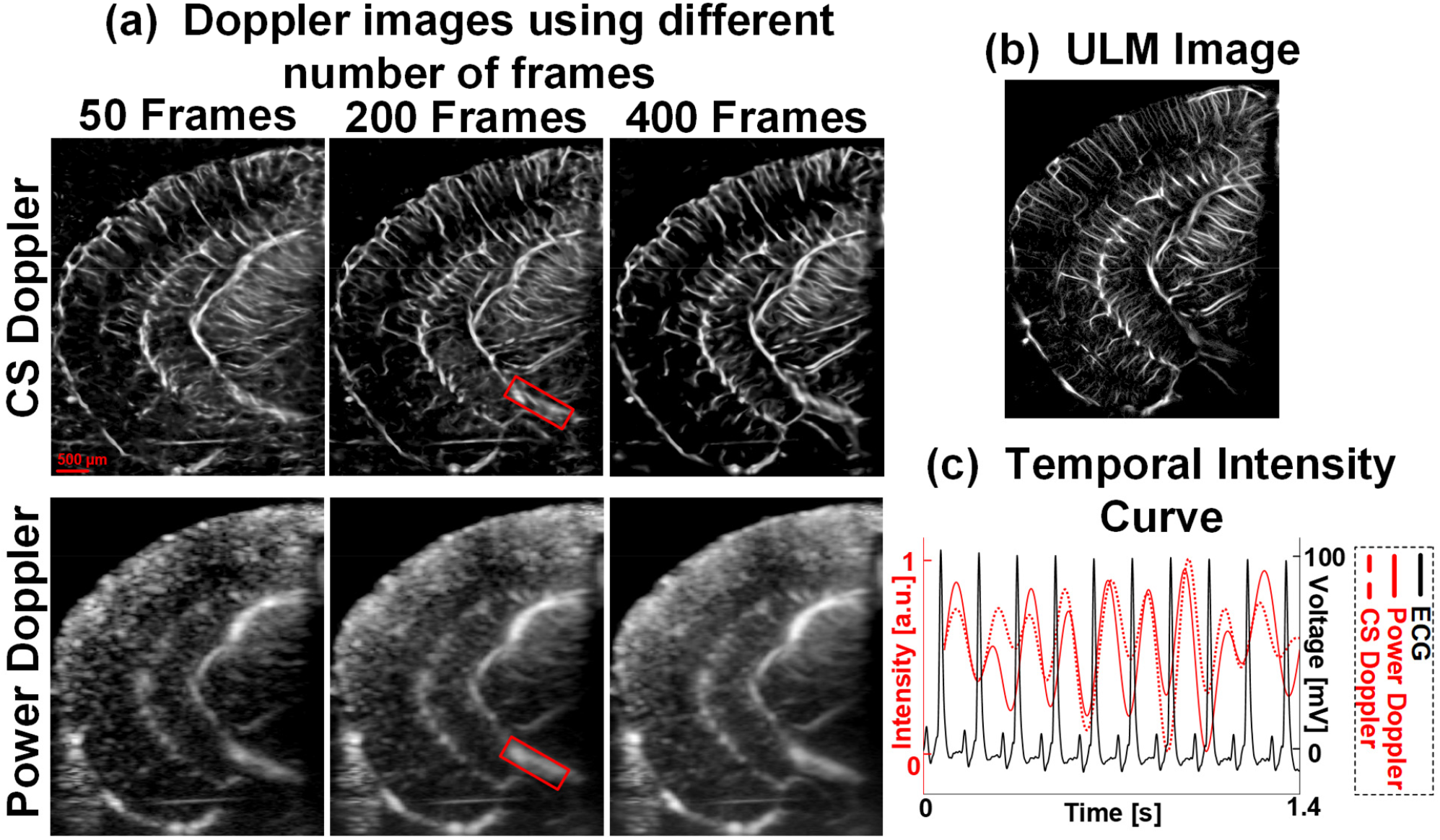
(a) CS Doppler and power Doppler images of the same mouse brain using different number of frames. (b) ULM image as ground truth. (c) Average intensity curve of the local region marked by the red rectangle in (a) along with the ECG signal. The temporal intensity curve was also demonstrated in the supplementary video 1.

### D. Demonstration of generalizing CS Doppler to other tissues not included in training datasets

Figure 6 shows the performance of CS Doppler in different types of tissues that were never seen by the deep generative network during training and validation. Both power Doppler and CS Doppler images were reconstructed using 0.4s of contrast-free data acquisition (400 frames). The CS Doppler images were reconstructed using the network trained with the L1 loss only. Interestingly, as shown in Fig. 6, CS Doppler could reconstruct most of the vessel structures with substantially improved spatial resolution in all the testing cases. This result is not surprising because although different imaging settings (e.g., frequency, transducer) and tissues were used for *in vivo* human and CAM imaging, the characteristics of the structure for smaller vessels are similar for different tissues. CS Doppler did struggle with some of the major vessels that have substantially different size distributions than those in the training dataset (indicated by the orange arrows in Fig. 6). It is expected that CS Doppler will gain in performance of reconstructing the larger vessels when data from these larger vessels are including in the training dataset.

**Fig. 6.**
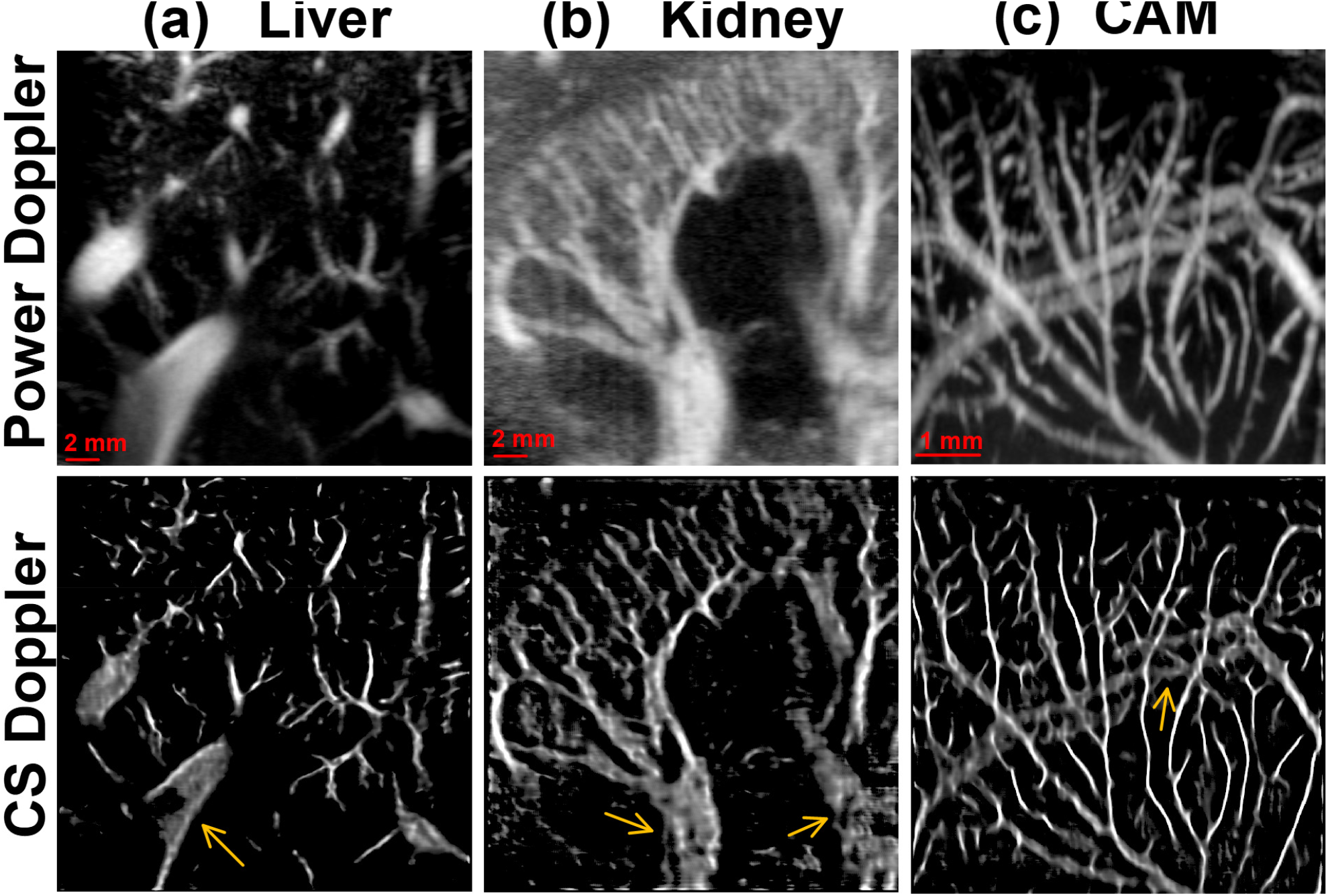
CS Doppler and power Doppler images of (a) a human liver (b) a human kidney (c) a chicken embryo CAM.

## IV. Discussion

This paper presented a contrast-free super-resolution Doppler (CS Doppler) imaging method that uses deep generative networks to reconstruct super-resolved microvessel structure from using spatiotemporal ultrasound signal. The paired ULM images were used as the ground truth during the training, validation and testing of the proposed generative network. Comprehensive assessments of the performance of CS Doppler were provided in this study. First, we compared the performance of CS Doppler using either with (e.g., raw spatiotemporal data) or without (e.g., accumulated power Doppler image) temporal information, and the result shows that temporal information was essential for recovering the microvessels with varying flow characteristics (Fig. 2). This finding is consistent with previous studies in optical and ultrasound super-resolution imaging (e.g., SOFI [21, 50]) where temporal blood flow information was utilized to break the diffraction limit of the imaging system.

Second, we used four different quantitative metrics (SSIM, PSNR, FWHM, MSE-NIP) to validate the performance of CS Doppler using different loss functions. Comparison studies with conventional Doppler and deconvolution-based Doppler were also conducted. Our results indicate that CS Doppler could improve the spatial resolution by an approximate factor of two when compared to conventional Doppler (Fig. 3(d)). More importantly, CS Doppler presented lower levels of hallucination when using the L1 loss, which is essential for super-resolution imaging in biomedical applications. In the study that compared different loss functions, the simplest loss function (e.g., L1 loss) demonstrated the best overall performance as indicated by the highest SSIM and MSE-NIP scores. Although the VGG loss and adversarial loss added more details and small vessel structures that delivered a higher PSNR, they did not provide good agreement with the reference standard ULM image, which indicates that these additional details and smaller vessel structures are more likely to be hallucinations [48-49]. This result is not surprising because the contrast-free data does not provide adequate information to support the full reconstruction of small vessel structures.

Third, we demonstrated that CS Doppler could preserve the temporal information of the power Doppler signal when using ECG as the common reference ground truth. As shown in Fig. 5 (c), CS Doppler presented a temporal blood flow profile that closely resembles the conventional power Doppler signal. This result is significant for applications where the intensity and temporal dynamics of the blood flow signal are utilized such as in tissue characterization and functional brain imaging.

Finally, we demonstrated the generalization ability of CS Doppler for different tissues that were not included in the training data. Although the training dataset only included ultrasound data acquired from mouse brain, CS Doppler still showed robust reconstruction results for human liver, human kidney and chicken embryo CAM (Fig. 6). This result suggests that CS Doppler is not limited to specific vessel structures associated with certain tissue types. As long as the testing data present similar vascular distributions as in the training data, CS Doppler will function and provide improved spatial resolution. Our results did indicate suboptimal performance in big vessels that were not included in the training data. To overcome this limitation, larger training dataset with wider vessel data distributions become necessary.

There are some limitations in this study. First, the contrast-enhanced data used for reconstructing the ULM ground truth images were acquired asynchronously with the contrast-free data. A small amount of tissue motion was present between different data acquisitions, which lead to misalignment between the contrast-free and contrast-enhanced data. Although motion correction was implemented [23] to correct for the mis-registration that was in-plane, out-of-plane motion remained. This misalignment between the contrast-free data and the ULM ground truth degrades the performance of the trained network. To address this issue, we used a combination of contrast-enhanced and contrast-free dataset for training. The contrast-enhanced data provides a strict correspondence between the flowing scatters and the microvessel structures for the network to learn, while the inclusion of the contrast-free data can adjust the network weights for better characterizing the distribution of the contrast-free dataset. This is a fundamental limitation of the proposed CS Doppler technique because it relies on paired ULM images for training. To address this issue, accelerated ULM imaging techniques such as CTSP [26], MUTE [28], and deep learning-based techniques [29-32] may be considered for fast imaging with less tissue motions. Nevertheless, based on the robust generalizability of the proposed method, one can also use transfer learning techniques [57] to adapt CS Doppler to different applications where only a small amount of application-specific ULM images is needed to fine-tune the network.

Another limitation of CS Doppler is that it is not expected to achieve the same resolution as ULM, especially for reconstructing small vessel structures below 20 µm (e.g., small arterioles, venules, and capillaries). Although CS Doppler uses ULM images as ground truth for training, contrast-free ultrasound signal does not have enough information to support the reconstruction of small vessels that are visible in ULM. However, compared with ULM, CS Doppler has a much higher temporal resolution and does not require contrast injection. These advantages make CS Doppler a more pragmatic and convenient imaging approach than ULM, which is significant for many preclinical and clinical applications.

## V. Conclusion

This paper presents a contrast-free super-resolution Doppler (CS Doppler) imaging method that uses deep generative networks to reconstruct the super-resolved microvessel image from the spatiotemporal ultrasound signal. The performance of the proposed method has been assessed by 1) measuring the quantitative metrics for different loss functions; 2) comparing the temporal intensity profile with conventional Doppler and electrocardiogram signal; 3) validating the results on different types of tissues beyond the training dataset. The results have shown that CS Doppler can robustly reconstruct the microvessel structure using contrast-free ultrasound signal with short acquisitions. This technique provides a practical solution for super-resolution microvessel imaging where fast imaging speed is essential.

## Reference

[1] O. M. Viessmann, R. J. Eckersley, K. Christensen-Jeffries, M. X. Tang, and C. Dunsby, “Acoustic super-resolution with ultrasound and microbubbles,” Phys. Med. Biol., vol. 58, no. 18, p. 6447, Sep. 2013.

[2] Desailly, Yann, Juliette Pierre, Olivier Couture, and Mickael Tanter. “Resolution limits of ultrafast ultrasound localization microscopy.” Physics in medicine & biology 60, no. 22 (2015): 8723.

[3] Y. Desailly, O. Couture, M. Fink, and M. Tanter, “Sono-activated ultrasound localization microscopy,” Appl. Phys. Lett., vol. 103, no. 17, 2013, Art. no. 174107.

[4] M. A. O’Reilly and K. Hynynen, “A super-resolution ultrasound method for brain vascular mapping,” Med. Phys., vol. 40, no. 11, 2013, Art. no. 110701.

[5] C. Errico et al., “Ultrafast ultrasound localization microscopy for deep super-resolution vascular imaging,” Nature, vol. 527, no. 7579, pp. 499–502, Nov. 2015.

[6] K. Christensen-Jeffries, R. J. Browning, M. X. Tang, C. Dunsby, and R. J. Eckersley, “In vivo acoustic super-resolution and super-resolved velocity mapping using microbubbles,” IEEE Trans. Med. Imag., vol. 34, no. 2, pp. 433–440, Feb. 2015.

[7] Lin, Fanglue, Sarah E. Shelton, David Espíndola, Juan D. Rojas, Gianmarco Pinton, and Paul A. Dayton. “3-D ultrasound localization microscopy for identifying microvascular morphology features of tumor angiogenesis at a resolution beyond the diffraction limit of conventional ultrasound.” Theranostics 7, no. 1 (2017): 196.

[8] Opacic, Tatjana, Stefanie Dencks, Benjamin Theek, Marion Piepenbrock, Dimitri Ackermann, Anne Rix, Twan Lammers et al. “Motion model ultrasound localization microscopy for preclinical and clinical multiparametric tumor characterization.” Nature communications 9, no. 1 (2018): 1–13.

[9] Harput, Sevan, Kirsten Christensen-Jeffries, Jemma Brown, Yuanwei Li, Katherine J. Williams, Alun H. Davies, Robert J. Eckersley, Christopher Dunsby, and Meng-Xing Tang. “Two-stage motion correction for super-resolution ultrasound imaging in human lower limb.” IEEE transactions on ultrasonics, ferroelectrics, and frequency control 65, no. 5 (2018): 803–814.

[10] [10] Zhu, Jiaqi, E. Rowland, Sevan Harput, Kai Riemer, Chee Hau Leow, Brett Clark, Karina Cox et al. “3D super-resolution ultrasound imaging of rabbit lymph node vasculature in vivo using microbubbles.” Radiological Society of North America, 2019.

[11] Lowerison, Matthew R., Wei Zhang, Xi Chen, Timothy M. Fan, and Pengfei Song. “Characterization of Anti-Angiogenic Chemo-Sensitization via Longitudinal Ultrasound Localization Microscopy in Colorectal Carcinoma Tumor Xenografts.” IEEE Transactions on Biomedical Engineering 69, no. 4 (2021): 1449–1460.

[12] Lowerison, Matthew R., Nathiya Vaithiyalingam Chandra Sekaran, Wei Zhang, Zhijie Dong, Xi Chen, Daniel A. Llano, and Pengfei Song. “Aging-related cerebral microvascular changes visualized using ultrasound localization microscopy in the living mouse.” Scientific reports 12, no. 1 (2022): 1–11.

[13] Huang, Chengwu, Wei Zhang, Ping Gong, U-Wai Lok, Shanshan Tang, Tinghui Yin, Xirui Zhang et al. “Super-resolution ultrasound localization microscopy based on a high frame-rate clinical ultrasound scanner: An in-human feasibility study.” Physics in Medicine & Biology 66, no. 8 (2021): 08NT01.

[14] Chavignon, Arthur, Baptiste Heiles, Vincent Hingot, Cyrille Orset, Denis Vivien, and Olivier Couture. “3D transcranial ultrasound localization microscopy in the rat brain with a multiplexed matrix probe.” IEEE Transactions on Biomedical Engineering 69, no. 7 (2021): 2132–2142.

[15] Demeulenaere, Oscar, Adrien Bertolo, Sophie Pezet, Nathalie Ialy-Radio, Bruno Osmanski, Clément Papadacci, Mickael Tanter, Thomas Deffieux, and Mathieu Pernot. “In vivo whole brain microvascular imaging in mice using transcranial 3D Ultrasound Localization Microscopy.” EBioMedicine 79 (2022): 103995.

[16] Demeulenaere, Oscar, Zulma Sandoval, Philippe Mateo, Alexandre Dizeux, Olivier Villemain, Romain Gallet, Bijan Ghaleh et al. “Coronary Flow Assessment Using 3-Dimensional Ultrafast Ultrasound Localization Microscopy.” JACC: Cardiovascular Imaging (2022).

[17] F. Lin, J. K. Tsuruta, J. D. Rojas, and P. A. Dayton, “Optimizing sensitivity of ultrasound contrast-enhanced super-resolution imaging by tailoring size distribution of microbubble contrast agent,” Ultrasound Med. Biol., vol. 43, no. 10, pp. 2488–2493, 2017.

[18] K. Christensen-Jeffries et al., “Microbubble axial localization errors in ultrasound super-resolution imaging,” IEEE Trans. Ultrason., Ferroelectr., Freq. Control, vol. 64, no. 11, pp. 1644–1654, Nov. 2017.

[19] P. Song, A. Manduca, J. D. Trzasko, R. E. Daigle, and S. Chen, “On the effects of spatial sampling quantization in super-resolution ultrasound microvessel imaging,” IEEE Trans. Ultrason., Ferroelectr., Freq. Control, vol. 65, no. 12, pp. 2264–2276, Dec. 2018.

[20] V. Hingot, C. Errico, M. Tanter, and O. Couture, “Subwavelength motion-correction for ultrafast ultrasound localization microscopy,” Ultrasonics, vol. 77, pp. 17–21, May 2017.

[21] A. Bar-Zion, C. Tremblay-Darveau, O. Solomon, D. Adam, and Y. C. Eldar, “Fast vascular ultrasound imaging with enhanced spatial resolution and background rejection,” IEEE Trans. Med. Imag., vol. 36, no. 1, pp. 169–180, Jan. 2017.

[22] A. Bar-Zion, O. Solomon, C. Tremblay-Darveau, D. Adam, and Y. C. Eldar, “SUSHI: Sparsity-based ultrasound super-resolution hemodynamic imaging,” IEEE Trans. Ultrason., Ferroelectr., Freq. Control, vol. 65, no. 12, pp. 2365–2380, Dec. 2018.

[23] P. Song et al., “Improved super-resolution ultrasound microvessel imaging with spatiotemporal nonlocal means filtering and bipartite graph-based microbubble tracking,” IEEE Trans. Ultrason., Ferroelectr., Freq. Control, vol. 65, no. 2, pp. 149–167, Feb. 2018.

[24] S. Tang et al., “Kalman filter-based microbubble tracking forrobust super-resolution ultrasound microvessel imaging,” IEEE Trans.Ultrason., Ferroelectr., Freq. Control, vol. 67, no. 9, pp. 1738–1751, Sep. 2020.

[25] C. Huang et al., “Short acquisition time super-resolution ultrasoundmicrovessel imaging via microbubble separation,” Sci. Rep., vol. 10, no. 1, pp. 1–13, Dec. 2020.

[26] Q. You et al., “Curvelet Transform-based Sparsity Promoting Algorithm for Fast Ultrasound Localization Microscopy,” in IEEE Transactions on Medical Imaging, doi: 10.1109/TMI.2022.3162839.

[27] Heiles, Baptiste, Arthur Chavignon, Vincent Hingot, Pauline Lopez, Eliott Teston, and Olivier Couture. “Performance benchmarking of microbubble-localization algorithms for ultrasound localization microscopy.” Nature Biomedical Engineering 6, no. 5 (2022): 605–616.

[28] Kim, Jihun, Mathew R. Lowerison, Nathiya V. Chandra Sekaran, Zhengchang Kou, Zhijie Dong, Michael L. Oelze, Daniel A. Llano, and Pengfei Song. “Improved Ultrasound Localization Microscopy Based on Microbubble Uncoupling via Transmit Excitation.” IEEE Transactions on Ultrasonics, Ferroelectrics, and Frequency Control 69, no. 3 (2022): 1041–1052.

[29] van Sloun, Ruud JG, Oren Solomon, Matthew Bruce, Zin Z. Khaing, Hessel Wijkstra, Yonina C. Eldar, and Massimo Mischi. “Super-resolution ultrasound localization microscopy through deep learning.” IEEE transactions on medical imaging 40, no. 3 (2020): 829–839.

[30] Lok, U-Wai, Chengwu Huang, Ping Gong, Shanshan Tang, Lulu Yang, Wei Zhang, Yohan Kim et al. “Fast super-resolution ultrasound microvessel imaging using spatiotemporal data with deep fully convolutional neural network.” Physics in Medicine & Biology 66, no. 7 (2021): 075005.

[31] Milecki, Léo, Jonathan Porée, Hatim Belgharbi, Chloé Bourquin, Rafat Damseh, Patrick Delafontaine-Martel, Frédéric Lesage, Maxime Gasse, and Jean Provost. “A deep learning framework for spatiotemporal ultrasound localization microscopy.” IEEE Transactions on Medical Imaging 40, no. 5 (2021): 1428–1437.

[32] Chen, Xi, Matthew Lowerison, Zhijie Dong, Nathiya Chandra Sekaran, Chengwu Huang, Shigao Chen, Timothy M. Fan, Daniel A. Llano, and Pengfei Song. “Localization free super-resolution microbubble velocimetry using a long short-term memory neural network.” bioRxiv (2021).

[33] V. Hingot, C. Errico, B. Heiles, L. Rahal, M. Tanter, and O. Couture, “Microvascular flow dictates the compromise between spatial resolution and acquisition time in ultrasound localization microscopy,” Sci. Rep., vol. 9, no. 1, pp. 1–10, Dec. 2019.

[34] M. R. Lowerison, C. Huang, Y. Kim, F. Lucien, S. Chen, and P. Song, “In vivo confocal imaging of fluorescently labeled microbubbles: Implications for ultrasound localization microscopy,” IEEE Trans. Ultrason., Ferroelectr., Freq. Control, vol. 67, no. 9, pp. 1811–1819, Sep. 2020.

[35] Christensen-Jeffries, Kirsten, Jemma Brown, Sevan Harput, Ge Zhang, Jiaqi Zhu, Meng-Xing Tang, Christopher Dunsby, and Robert J. Eckersley. “Poisson statistical model of ultrasound super-resolution imaging acquisition time.” IEEE transactions on ultrasonics, ferroelectrics, and frequency control 66, no. 7 (2019): 1246–1254.

[36] Demené, Charlie, Thomas Deffieux, Mathieu Pernot, Bruno-Félix Osmanski, Valérie Biran, Jean-Luc Gennisson, Lim-Anna Sieu et al. “Spatiotemporal clutter filtering of ultrafast ultrasound data highly increases Doppler and fUltrasound sensitivity.” IEEE transactions on medical imaging 34, no. 11 (2015): 2271–2285.

[37] Baranger, Jérôme, Bastien Arnal, Fabienne Perren, Olivier Baud, Mickael Tanter, and Charlie Demené. “Adaptive spatiotemporal SVD clutter filtering for ultrafast Doppler imaging using similarity of spatial singular vectors.” IEEE transactions on medical imaging 37, no. 7 (2018): 1574–1586.

[38] Song, Pengfei, Armando Manduca, Joshua D. Trzasko, and Shigao Chen. “Ultrasound small vessel imaging with block-wise adaptive local clutter filtering.” IEEE transactions on medical imaging 36, no. 1 (2016): 251–262.

[39] Bar-Zion, Avinoam, Oren Solomon, Claire Rabut, David Maresca, Yonina C. Eldar, and Mikhail G. Shapiro. “Doppler Slicing for Ultrasound Super-Resolution Without Contrast Agents.” bioRxiv (2021).

[40] Lok, U-Wai, Joshua D. Trzasko, Chengwu Huang, Shanshan Tang, Ping Gong, Yohan Kim, Fabrice Lucien, Matthew R. Lowerison, Pengfei Song, and Shigao Chen. “Improved ultrasound microvessel imaging using deconvolution with total variation regularization.” Ultrasound in Medicine & Biology 47, no. 4 (2021): 1089–1098.

[41] Jensen, Jørgen Arendt, Mikkel Schou, Sofie Bech Andersen, Stinne Byrholdt Søgaard, Charlotte Mehlin Sørensen, Michael Bachmann Nielsen, Carsten Gundlach et al. “Fast super resolution ultrasound imaging using the erythrocytes.” In Medical Imaging 2022: Ultrasonic Imaging and Tomography, vol. 12038, pp. 79-84. SPIE, 2022.

[42] Park, Jun Hong, Woorak Choi, Gun Young Yoon, and Sang Joon Lee. “Deep learning-based super-resolution ultrasound speckle tracking velocimetry.” Ultrasound in Medicine & Biology 46, no. 3 (2020): 598–609.

[43] Kim, Jongbeom, Gyuwon Kim, Lei Li, Pengfei Zhang, Jin Young Kim, Yeonggeun Kim, Hyung Ham Kim, Lihong V. Wang, Seungchul Lee, and Chulhong Kim. “Deep learning acceleration of multiscale superresolution localization photoacoustic imaging.” Light: Science & Applications 11, no. 1 (2022): 1–12.

[44] Ouyang, Wei, Andrey Aristov, Mickaël Lelek, Xian Hao, and Christophe Zimmer. “Deep learning massively accelerates super-resolution localization microscopy.” Nature biotechnology 36, no. 5 (2018): 460–468.

[45] Ledig, Christian, Lucas Theis, Ferenc Huszár, Jose Caballero, Andrew Cunningham, Alejandro Acosta, Andrew Aitken et al. “Photo-realistic single image super-resolution using a generative adversarial network.” In Proceedings of the IEEE conference on computer vision and pattern recognition, pp. 4681-4690. 2017.

[46] Wang, Xintao, Ke Yu, Shixiang Wu, Jinjin Gu, Yihao Liu, Chao Dong, Yu Qiao, and Chen Change Loy. “Esrgan: Enhanced super-resolution generative adversarial networks.” In Proceedings of the European conference on computer vision (ECCV) workshops, pp. 0-0. 2018.

[47] Chen, Yuhua, Feng Shi, Anthony G. Christodoulou, Yibin Xie, Zhengwei Zhou, and Debiao Li. “Efficient and accurate MRI super-resolution using a generative adversarial network and 3D multi-level densely connected network.” In International Conference on Medical Image Computing and Computer-Assisted Intervention, pp. 91–99. Springer, Cham, 2018.

[48] Bhadra, Sayantan, Varun A. Kelkar, Frank J. Brooks, and Mark A. Anastasio. “On hallucinations in tomographic image reconstruction.” IEEE transactions on medical imaging 40, no. 11 (2021): 3249–3260.

[49] Kelkar, Varun A., Dimitrios S. Gotsis, Frank J. Brooks, Prabhat KC, Kyle J. Myers, Rongping Zeng, and Mark A. Anastasio. “Assessing the ability of generative adversarial networks to learn canonical medical image statistics.” arXiv preprint 2204.12007 (2022).

[50] Dertinger, Thomas, Ryan Colyer, Gopal Iyer, Shimon Weiss, and Jörg Enderlein. “Fast, background-free, 3D super-resolution optical fluctuation imaging (SOFI).” Proceedings of the National Academy of Sciences 106, no. 52 (2009): 22287–22292.

[51] Simonyan, Karen, and Andrew Zisserman. “Very deep convolutional networks for large-scale image recognition.” arXiv preprint 1409.1556 (2014).

[52] Goodfellow, Ian, Jean Pouget-Abadie, Mehdi Mirza, Bing Xu, David Warde-Farley, Sherjil Ozair, Aaron Courville, and Yoshua Bengio. “Generative adversarial nets.” Advances in neural information processing systems 27 (2014).

[53] Dosovitskiy, Alexey, and Thomas Brox. “Generating images with perceptual similarity metrics based on deep networks.” Advances in neural information processing systems29 (2016).

[54] Kingma, Diederik P., and Jimmy Ba. “Adam: A method for stochastic optimization.” arXiv preprint 1412.6980 (2014).

[55] Arjovsky, Martin, Soumith Chintala, and Léon Bottou. “Wasserstein generative adversarial networks.” In International conference on machine learning, pp. 214-223. PMLR, 2017.

[56] Wang, Zhou, Alan C. Bovik, Hamid R. Sheikh, and Eero P. Simoncelli. “Image quality assessment: from error visibility to structural similarity.” IEEE transactions on image processing 13, no. 4 (2004): 600–612.

[57] Pan, Sinno Jialin, and Qiang Yang. “A survey on transfer learning.” IEEE Transactions on knowledge and data engineering 22, no. 10 (2009): 1345–1359.

